# A negative autoregulation network motif is required for synchronized *Myxococcus xanthus* development

**DOI:** 10.1101/738716

**Authors:** Patrick T. McLaughlin, Penelope I. Higgs

## Abstract

Transcription factor autoregulation is a simple network motif (recurring circuit) built into genetic regulatory networks that direct cell behavior. Negative autoregulation (NAR) network motifs are particularly abundant in bacteria and provide specific functions, such as buffering against transcriptional noise. Here, we investigate the phenotypic consequence of perturbing NAR of a major transcription factor, MrpC, that controls the multicellular development program of the bacterium *Myxococcus xanthus*. Launch of the developmental program directs certain cells in the population to first aggregate into haystack-shaped mounds, and then to differentiate into environmentally resistant spores to form mature fruiting bodies. Perturbation of MrpC NAR causes a striking phenotype in which cells lose synchronized transition from aggregation to sporulation. Instead, some cells abruptly exit aggregation centers and remain locked in a cohesive swarming state, while the remaining cells transition to spores inside residual fruiting bodies. As predicted, disruption of MrpC NAR led to an increased and broadened population distribution of *mrpC* expression. Examination of MrpC levels in developmental subpopulations during *in situ* development demonstrated cells locked in the swarms contained intermediate MrpC levels insufficient to promote sporulation. These results suggest an inherent property of NAR motifs that function in multicellular developmental programs is to facilitate synchronized responses.

**Significance Statement:** All organisms use regulatory networks for cellular homeostasis, mediating appropriate responses to environmental changes, or to direct animal development. Understanding how the basic building blocks (motifs) of regulatory networks contribute to these processes is essential to mitigate the effects of mutations in regulatory networks (*i.e.* cancers) or to synthesize beneficial organisms. In this study, we demonstrate that a common regulatory motif, a transcription factor that represses its own expression, helps synchronize cells that engage in collective behaviors.

## Introduction

Genetic regulatory networks (GRNs) can be thought of as algorithms that direct organisms to perform simple single-cell functions, such as appropriate response to environmental changes (1), or more complex multicellular programs such as embryogenesis (2). GRNs are built from recurring simple regulatory circuits (network motifs) which each perform distinct functions (1). The simplest of these network motifs is autoregulation, in which a transcription factor enhances (positive autoregulation; PAR) or represses (negative autoregulation; NAR) its own expression. NAR is a particularly abundant network motif (3, 4). Theoretical and experimental data, derived mostly from isolated circuits, have demonstrated NAR network motifs buffer against transcriptional noise, speed up response times, and increase the input dynamic range of a circuit (4-6). Most of these well-described functions have been investigated in single celled organisms or in synthetic systems, whereas few examples of the phenotypic consequences of network motifs in natural multicellular systems are available.

*Myxococcus xanthus* is a “social” bacterium with a life cycle that is highly dependent on collective behaviors (7). During vegetative growth, large groups of *M*. *xanthus* cells swarm over solid surfaces in search of prey on which they cooperatively predate (8, 9). Under nutrient limited conditions, *M*. *xanthus* enters a dedicated developmental program during which cells form a specialized biofilm and segregate into distinct cell fates (10). Some cells are induced to swarm into aggregation centers to produce mounds of approximately 10^5^ cells. Once inside aggregation centers, individual cells slow down and stop moving (11), which prevents them from leaving the aggregation center. These cells are then induced to differentiate into environmentally resistant spores, producing mature multicellular fruiting bodies. Other cells within the developing population remain outside of the fruiting bodies and differentiate into peripheral rods, which are thought of as a persister-like state (12, 13). For development to be effective, cells in the population must coordinate their behavior both in time and space. If sporulation were to occur before completion of aggregation, then the benefits conferred by a multicellular fruiting body structure, such as enhanced resistance to environmental stresses or group dispersal, would be lost (14).

A GRN controlling the *M*. *xanthus* developmental program has been largely defined [reviewed in (15)]. A key player near the top of the developmental GRN is MrpC, a CRP/Fnr superfamily transcriptional regulator that regulates hundreds of developmental genes (16, 17). Threshold levels of MrpC drive distinct stages of development: low levels are associated with induction of the aggregation onset, higher levels are associated with commitment to sporulation (13, 18-20). Peripheral rods contain low levels of MrpC (13).

MrpC is under complex regulation. Shortly after cells sense nutrient limitation, *mrpC* expression is upregulated by MrpB, a bacterial enhancer binding protein (bEBP) (16). MrpB binds to two upstream activating sequences (UAS1 and UAS2) 182 bp from the *mrpC* start codon and is proposed to stimulate *mrpC* expression from a sigma^54^-dependent promoter (16, 21). Early during the developmental program, gradual accumulation of MrpC is achieved because the Esp signaling system induces turnover of MrpC (22, 23). Several additional post-transcriptional events modulate MrpC accumulation (and therefore progression through development) in response to changing environmental conditions (18, 19, 24). A long-standing model suggested MrpC acted as positive autoregulator, leading to the attractive speculation that bifurcation of MrpC production (a known feature of positive autoregulation motifs (25)) could drive cell fate segregation into either aggregation centers or peripheral rods [summarized in (15)]. Recently, however, we demonstrated that MrpC functions instead as a negative autoregulator by competing with its transcriptional activator, MrpB, for overlapping binding sites in the *mrpC* promoter region (21).

Here, we set out to understand the role of the MrpC NAR network motif in the natural GRN driving the *M*. *xanthus* developmental program. We demonstrate that perturbation of MrpC NAR leads to early and over-production of MrpC associated with premature aggregation onset, reduced fruiting body organization, and, unexpectedly, reduced sporulation efficiency. Using a recently developed submerged culture developmental filming method (26), we observed perturbed MrpC NAR resulted in asynchronous development: after formation of aggregates, some cells suddenly exited aggregation centers as fast-moving swarms, while other cells remained in stationary fruiting bodies. Deep convolution neural network analyses indicated these developmental swarms displayed trajectories and velocities that were distinct from cells in the mobile aggregate phase. Analysis of single cell *mrpC* expression demonstrated that perturbed MrpC NAR resulted in increased levels and cell-to-cell variability in *mrpC* expression. *In situ* analysis of single cell MrpC accumulation suggested that the developmental swarms exhibited intermediate MrpC levels. Together, these data indicate MrpC NAR functions to decrease cell-to-cell variability, which is crucial for synchronized transition from cells in the aggregation state to sporulation state. This study provides a clear example of the function of NAR network motifs in GRNs directing multicellular development.

## Results

### *mrpC* Negative Autoregulation Requires Four Distinct MrpC Binding Sites

*mrpC* expression is subject to NAR due at least in part to competition between MrpC and its transcriptional activator, MrpB, for overlapping binding sites (21) (Fig. S1A). MrpC binds to two additional sequences within its promoter (termed BS5 and BS1; Fig. S1A) (21). To examine whether BS5 and BS1 contributed to MrpC NAR, we analyzed *mrpC* expression reporter constructs bearing the wild type promoter or mutant promoters bearing perturbation of BS1 (BS1*), deletion of BS5 (ΔBS5), or both (ΔBS5 BS1*). BS1* was generated by substituting the TGT consensus resides to GAA which completely disrupts MrpC binding to BS1 *in vitro* (21)(Fig. S1B). Each reporter construct was inserted into the Mx8 phage attachment site (*attB*) of wild type *M. xanthus* strain DZ2, and developmental assays indicated all resulting strains displayed wild-type developmental phenotypes (data not shown). Analysis of the reporter during development demonstrated that deletion of BS5 resulted in a 2.53 ± 0.04-fold increase in mCherry fluorescence compared to the wild type at 48 h, and the BS1* mutation resulted in a 1.8 ± 0.1-fold increase relative to wild type (Fig. S1C). The reporter bearing the BS1* ΔBS5 double disruption yielded a 3.3 ± 0.2 and 1.29 ± 0.09-fold increase in mCherry fluorescence compared to the parent and ΔBS5 reporters, respectively (Fig. 1A and S1C). No significant mCherry fluorescence was detected from the P_BS5-3_-mCh reporter which lacks the MrpB binding sites (UAS1 and 2/BS3 and 4) (21)(Fig. S1C). Together, these results suggest that both BS1 and BS5 are necessary for appropriate NAR and play additive roles. The observation that reporter expression in the Δ*mrpC* strain is dramatically higher early during development (Fig. S1D) suggests that MrpC binding to BS3/4 (hindering MrpB binding at UAS1/2) also plays a role in appropriate NAR. We are unable to generate mutations in BS3/4 which wouldn’t also perturb MrpB-dependent activation of *mrpC* expression from UAS1/2 (21).

**Figure 1.**
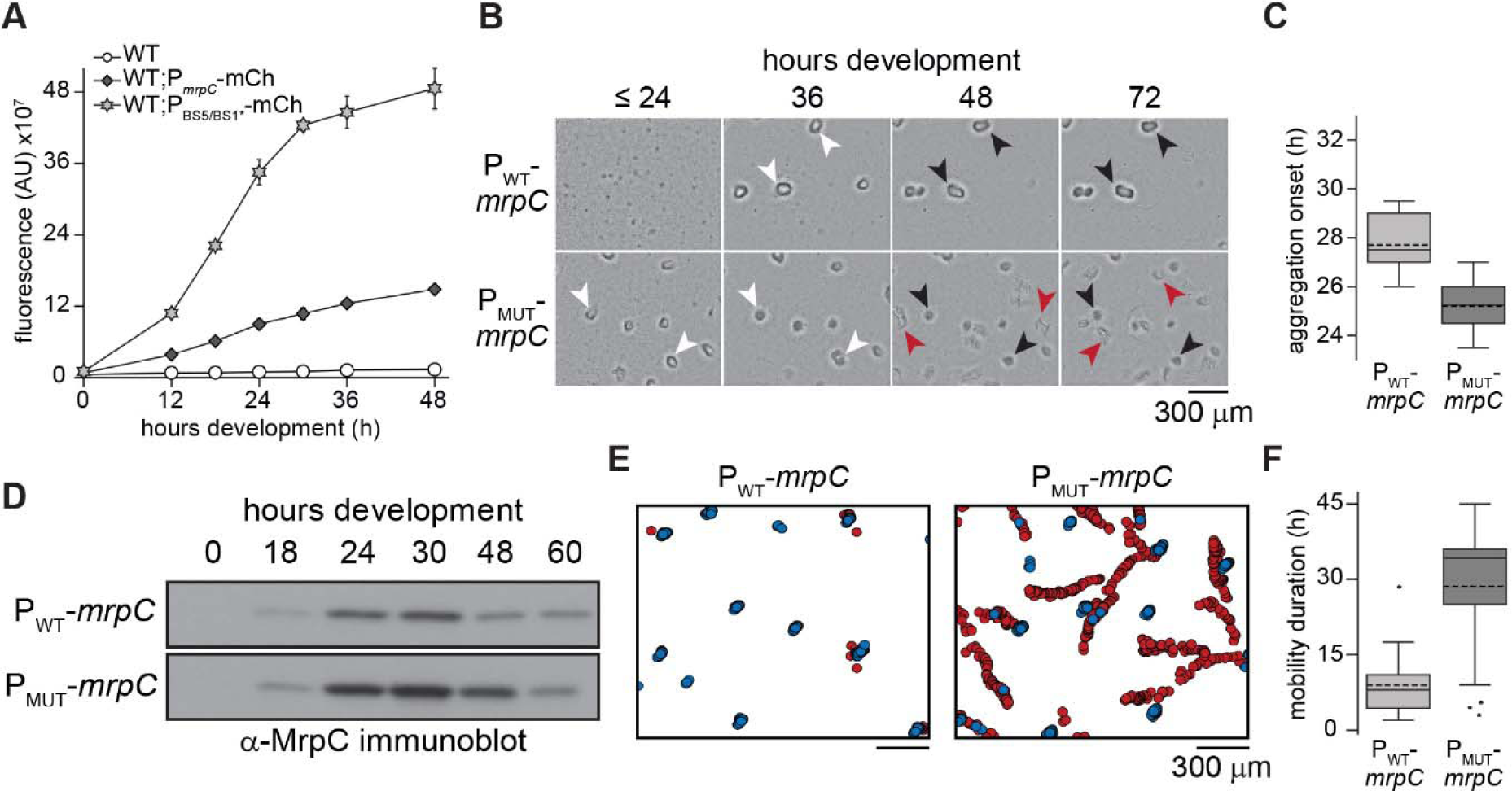
Perturbation of MrpC negative autoregulation (NAR) leads to asynchronous development. (A) *mrpC* expression is increased when binding sites (BS) 5 and 1 are disrupted. Analysis of *mrpC* expression with an mCherry reporter expressed from the wild type *mrpC* (P_mrpC_-*mCh*) or perturbed MrpC NAR (P_BS5/BS1*_-*mCh mrpC*) promoters. Reporters were integrated into wild type cells and mCherry fluorescence was recorded during development under submerged culture. (B) Perturbation of MrpC NAR caused early aggregation and disrupted fruiting bodies. The developmental phenotype observed from Δ**mrpC** cells expressing *mrpC* from its wild type-(P_WT_-*mrpC*) or P_BS5/BS1-_ (P_MUT_-*mrpC*) promoters. White arrows: aggregates of ∼10^5^ cells; black arrows: immobile fruiting bodies; red arrows: metastatic swarms. (C, F) Distribution of developmental times at which aggregation centers were first observed (aggregation onset; C), or durations of time that aggregates (P_WT_-*mrpC*) or metastatic swarms (P_MUT_-*mrpC*) traveled (mobility duration; F). Strains were developed under submerged culture in 96 well plates and imaged every 30 min for 72 hours. n=15 from three independent biological replicates. Solid line: median; dashed line: mean. (D) Anti-MrpC immunoblot of lysates prepared from P_WT_-*mrpC* or P_MUT_-*mrpC* cells developing under submerged culture. (E) Identification and tracking of aggregates by a DeepLabCut trained neural network. Stationary aggregates (blue) and metastatic swarms (red) identified from movies of cells developing under submerged culture as in C. Data from one representative assay is shown.

### Perturbed MrpC NAR Leads to Early Aggregation and Reduced Sporulation

To examine how perturbation of MrpC NAR affects *M. xanthus* development, we next generated constructs in which *mrpC* was driven from its wild type promoter (P_WT_) or from the P_BS5/BS1*_ promoter (P BS5/BS1*). These constructs were integrated in the Δ*mrpC* background at the *attB* site, producing Δ*mrpC attB*::P_WT_-*mrpC* (parent) and Δ*mrpC attB*::P _BS5/BS1*_-*mrpC* strains (hereafter termed P_WT_-*mrpC* and P_MUT_-*mrpC*, respectively). The developmental phenotype of these two strains was compared to the Δ*mrpC* and wild type background strains. Under submerged culture developmental conditions, the Δ*mrpC* strain failed to produce aggregates, whereas the wildtype and P_WT_-*mrpC* strains produced visible aggregation centers between 24 to 36 h that darkened by 72 h post-starvation (Fig. S2A and 1B), indicating exogenously expressed *mrpC* restores wild type aggregation. In contrast, the P_MUT_-*mrpC* strain produced aggregates at least 6 h earlier than the parent strain (Fig. S2A and 1B). Furthermore, while the P_MUT_-*mrpC* aggregates appeared similar to the parent at 36 h, they subsequently failed to appropriately darken, and by 48-72 h became more disorganized than the parent and wild-type strains (Fig. 1B).

To examine sporulation levels in these strains, heat- and sonication-resistant spores were enumerated at 48, 72, and 120 h. After 120 h of development, the wild type had produced 3.1 ± 0.7 × 10^7^ spores (recorded as 100 ± 23%), while no spores could be detected from Δ*mrpC* mutant (≤ 0.07% wild type)(Fig. S2B). The P_WT_-*mrpC* strain produced 70 ± 16% of the wild type levels, suggesting exogenous expression of *mrpC* did not fully complement with respect to sporulation efficiency (Fig. S2B). The P_MUT_-*mrpC* strain, however, exhibited a striking reduction in sporulation corresponding to 30 ± 17% of the resistant spores produced by the P_WT_-*mrpC* strain (Fig. S2B). To determine if there was an inherent defect in the core sporulation program in the P_MUT_-*mrpC* mutant, we examined the number of heat- and sonication-resistant spores produced after artificial chemical induction of sporulation in liquid cultures which bypasses the requirement for aggregation (27). The P_MUT_-*mrpC* mutant produced a similar number of chemical-induced spores as the wild type, whereas the Δ*mrpC* mutant failed to produce any spores (Fig. S2C). These results suggested that the inefficient sporulation observed by the P_MUT_-*mrpC* strain during starvation-induced development was not due to failure to execute spore differentiation *per se*.

Finally, to examine how the observed phenotypes correlated with total MrpC levels, we prepared lysates from P_WT_-*mrpC* or P_MUT_-*mrpC* strains at intervals between 0 – 60 hours of development and subjected them to anti-MrpC immunoblot. In the P_WT_-*mrpC* strain, MrpC was absent at the onset of development (t = 0), increased between 18 – 30 h of development, and then subsequently decreased after the onset of sporulation (Fig. 1D and S2B). Relative to the P_WT_-*mrpC* strain, levels of MrpC in the P_MUT_-*mrpC* strain were 2-3-fold higher between 18-48 h, and eventually decreased to P_WT_-*mrpC* levels by 60 h (Figs. 1D and S4B). This pattern of MrpC accumulation in the two strains was similar to *mrpC* expression levels when the longevity of the mCherry reporter was accounted for by plotting the derivative of the mCherry fluorescence (Fig. S4A). Thus, elevated MrpC levels at 24 hours likely explained the early aggregation onset observed in the P_MUT_-*mrpC* strain (Fig 1B). However, the reduced sporulation efficiency observed by this strain was surprising given MrpC levels were similar at 60 h of development when sporulation levels were reduced compared to the parent (Fig. S2B). Together, these results suggested that perturbing MrpC NAR resulted in an uncoupling between completion of aggregation and induction of sporulation.

### Perturbation of MrpC NAR Leads to Asynchronous Development

Movies of *M. xanthus* development have demonstrated that prior to the onset of sporulation, aggregates are surprisingly dynamic (26, 28). Initial aggregates often dissolve or coalesce, and even mature aggregates can be mobile prior to transition to stationary spore-filled fruiting bodies (26). To examine how MrpC NAR affected these transient behaviors, the wild type, P_WT_-*mrpC*, and P_MUT_-*mrpC* strains were induced to develop under submerged culture conditions, imaged every 30 min from 0 – 72 h in an automated plate reader, and the images assembled into movies. For each movie, the time of onset of aggregation, the number of initial versus mature aggregates, and the duration of mature aggregate mobility was recorded (Movie 1 and 2). These analyses demonstrated aggregation onset in the P_MUT_-*mrpC* strain assays was 3 hours earlier than the wild type and P_WT_-*mrpC* strains (25 ± 1 vs. 28 ± 1 and 28 ± 1 h post-starvation, respectively) (Figs. 1C and S2D)(Table S1).

After mature aggregates were formed, remarkable differences in their behavior was observed. In the wild type strain, very few mobile aggregates were observed (5 ± 12 %) and the duration of mobility was short (2 ± 1 h) (Table S1). In the P_WT_-*mrpC* strain, 40 ± 30 % of the aggregates were mobile for 8 ± 6 h (Fig. 1E-F, Table S1)(Movie 1). As the endogenous *mrpC* promoter region is still intact in the Δ*mrpC* background, we speculated that the differences between the wild type and P_WT_-*mrpC* (i.e. Δ*mrpC attB*::P_WT_-*mrpC*) strains may result from two copies of the *mrpC* promoter region in this strain which may dilute the NAR activity of the available MrpC. Remarkably, however, in 80 ± 20 % of the aggregates produced in the P_MUT_-*mrpC* strain, a large proportion of cells suddenly exited one side of the aggregate, leaving other cells behind as darkened shallow fruiting bodies (Movie 2). The cells that exited the aggregate collectively migrated throughout the plate; we referred to these cells as a metastatic swarm. Most of these swarms (70 %; 19/27) were still actively moving by the end of the filming period at 72 h. These observations explained both the disorganized appearance of the late aggregates (Fig. 1B) and the reduced sporulation efficiencies (Fig. S2B) observed in the P_MUT_-*mrpC* strain.

On average, all three strains produced similar maximum numbers of initial aggregates (between 6-8) and final fruiting bodies (between 5-6) (Table S1). However, in the P_MUT_-*mrpC* strain, the overall number of aggregates that transitioned into fruiting bodies (60 ± 10%) was significantly reduced (*p* = 0.031) compared to the wild type and P_WT_-*mrpC* strains (80 ± 16 % and 80 ± 19 %, respectively) (Table S1, Fig. S3B). Interestingly, although aggregation onset in the P_MUT_-*mrpC* cells was at least 2 h earlier than the other strains, there was no significant difference in the time at which fully formed aggregates began to move in the P_MUT_-*mrpC* (37 ± 4 h) relative to the P_WT_-*mrpC* cells (37 ± 3 h) (Table S1, Fig. S3A). Thus, the interval between formation of aggregates and onset of aggregate mobility in the P_WT_-*mrpC* and P_MUT_-*mrpC* was 7 ± 3, and 11 ± 3 h, respectively.

To better compare the characteristics of the individual entities produced by the P_MUT_-*mrpC* and P_WT_-*mrpC* mobile strains, we employed the DeepLabCut deep convolutional neural network (29) to analyze developing strains. The neural network was trained on manually labelled P_WT_-*mrpC* or P_MUT_-*mrpC* developmental images with a 50-layer residual network (ResNet-50) for 340,000 iterations resulting in a training and test error of 1.62 and 6.66 pixels, respectively (30). The trained neural network was then used to analyze fifteen videos of each strain (3 independent biological replicates each with 5 technical replicates). First, the neural network assigned non-mobile aggregates a median speed of 0.22 μm/min (IQR: 0.12 – 0.40 μm/min), which was attributed to an artifact from slight shifts in the position of the automated plate reader camera between time points (data not shown). In the P_WT_-*mrpC* strain, the mobile aggregates travelled at an average speed of 0.3 μm/min (Fig. S3F), and movement was primarily radial (Fig. 1E and S3C) and the average net displacement was within two aggregate diameters of the starting point (Fig. 1E and S3E). In the P_MUT_-*mrpC* strain, metastatic swarms travelled at an average speed of 0.6 μm/min (Fig. S3F) and movement involved included long runs, sharp turns, and/or repeated reversals (Fig. 1E and S3D) such that average total displacement was 1500 ± 400 μm (Fig. S3E). Unlike with the P_WT_-*mrpC* strain, almost all mobile aggregates left a shallow immobile fruiting body behind (Fig. 1B and E). Thus, metastatic swarms displayed speed and trajectory characteristics that were significantly different from the P_WT_-*mrpC* mobile aggregates, suggesting they did not result from simply increasing the duration of the normal aggregate mobility phase.

### MrpC NAR Dampens Cell-to-cell Variability in *mrpC* Expression

Theoretical and experimental analyses have suggested that NAR motifs function to decrease cell-to-cell variability in gene expression, ensuring that expression is homogenous within a population (5). Since the P_MUT_-*mrpC* strain appeared to produce subpopulations of cells in different developmental states (i.e. metastatic swarms and stationary fruiting bodies; Fig. 1E), we hypothesized that MrpC-mediated NAR may be functioning to limit heterogeneity of *mrpC* expression thus ensuring a coordinated developmental process.

To examine population variation in *mrpC* expression, we measured mCherry production from individual wild-type cells bearing the P_WT_-mCh or P_MUT_-mCh reporter during *in situ* development under submerged culture. For normalization purposes, these cells also contained a construct that expressed mNeonGreen from an inducible promoter integrated at a second genomic site (1.38kb::P_vanillate_-*mNeonGreen*). These double labeled strains were each diluted 1:19 into a markerless wild type and single cell fluorescence was recorded using confocal laser scanning microscopy (CLSM). For each cell, mCherry fluorescence was normalized to mNeonGreen fluorescence, and the distribution of red-to-green (RG) ratios from individual cells from three independent biological experiments was plotted. Variability in single cell RG ratios amongst the population was calculated using the coefficient of variation (CV).

In pre-aggregating cells (24 h development) with the wild type reporter, a mean RG ratio of 0.38 ± 0.09 was observed with a narrow distribution in values (Fig. 2A) that corresponded to a CV of 23.4 ± 0.3% (Fig. 2D). Cells with the P_MUT_-mCh reporter displayed both 3-fold increased mean reporter expression (RG ratio of 1.2 ± 0.4) and a significantly increased spread in distribution of expression (CV 32 ± 2 %)(Fig. 2A and D). Similarly, in fruiting bodies, cells bearing the P_MUT_-mCh reporter displayed 2.5- and 1.3-fold increased mean RG ratio and CV compared to the P_WT_-mCh fruiting body cells, respectively (Fig. 2B and D). This trend was also observed in the peripheral rods (mean RG ratio and CV of 3- and 1.3-fold higher than the P_WT_-mCh reporter, respectively). These results indicated that MrpC-mediated NAR functions to not only limit the level of expression, but also to limit the cell-to-cell variability in expression. Additionally, these results suggested the increase in variability upon perturbation of NAR was not subpopulation specific, indicating that MrpC-mediated NAR was functioning throughout the entire population to limit expression heterogeneity.

**Figure 2.**
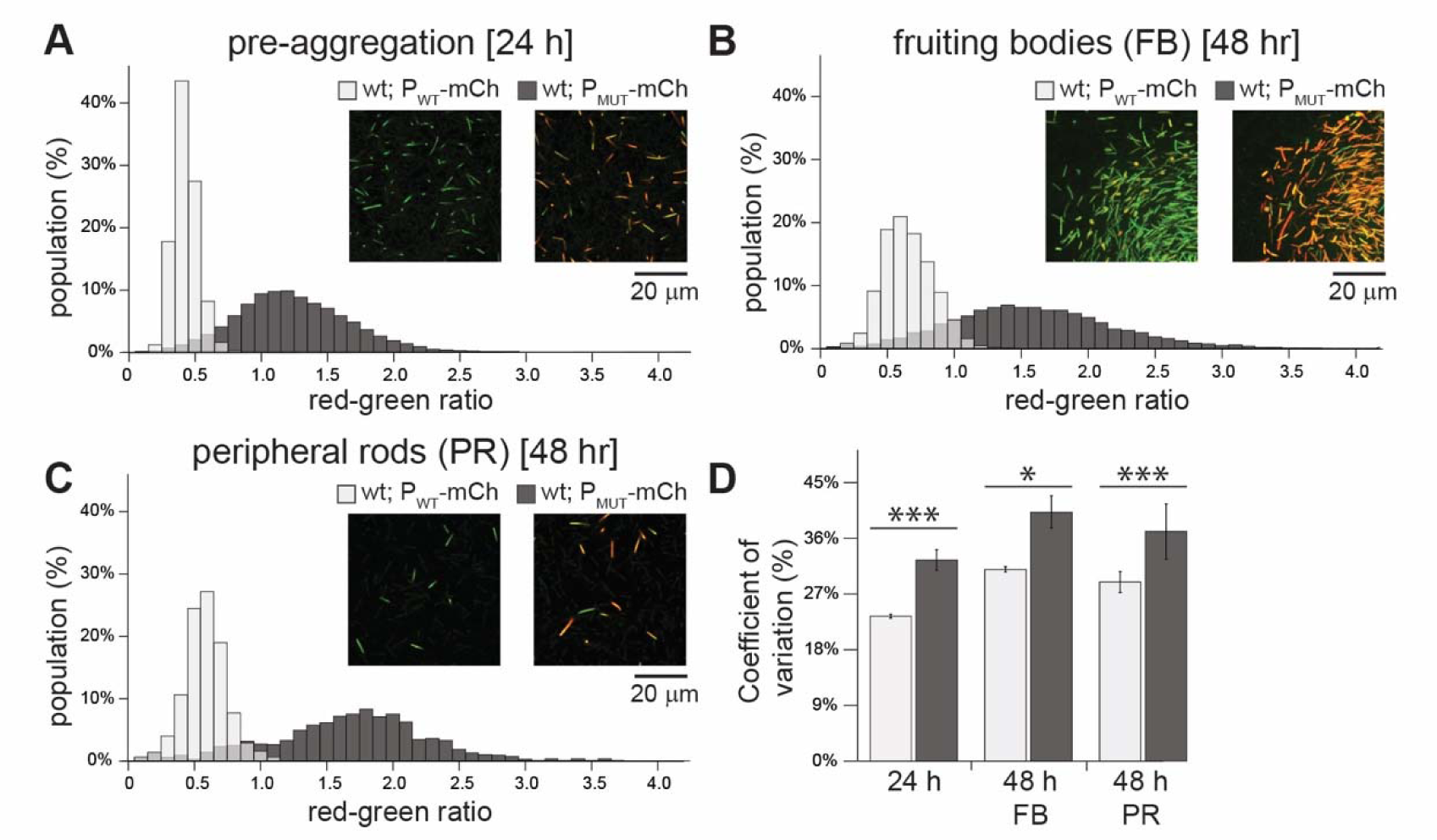
Perturbation of MrpC NAR increases the level and population-wide variability of *mrpC* expression. (A-C) Histogram of *mrpC* reporter expression in pre-aggregation (A), fruiting body (B), or peripheral rod populations (C). Wild-type cells expressing mCherry from either the wild-type *mrpC* promoter (P_mrpC_-*mCh*; P_WT_-*mCh*) or perturbed NAR *mrpC* promoter (P_BS5/BS1*_-*mCh*; P_MUT_-*mCh*), and mNeonGreen from a constitutive promoter (P_vanillate_-*mNG*). Strains were developed under submerged culture and imaged by confocal microscopy. mCherry (red) and mNeonGreen fluorescence was measured for individual cells and the red-green ratio calculated. Ratios were binned and the percent of the population with the indicated red-green ratio displayed by histogram. For the P_WT_-mCh strain, n = 8064 (A), 14,298 (B), and 933 (C) cells. For the P_MUT_-mCh strain, n = 5835 (A), 11,198 (B), and 850 (C) cells. Representative images showing mCherry and mNeonGreen fluorescence over-layed from each population in the indicated strains is shown. Note: the edge of a fruiting body is shown in B. (D) Average coefficient of variation and associated standard deviation was calculated from three independent biological replicates. Light gray: P_WT_-*mCh*; dark gray: P_MUT_-*mCh*. Data was analyzed for statistically significant differences using the unequal variance T-test. *: *p* < 0.05; *** *p* < 0.001.

### Developmental Swarms Possess an Intermediate level of MrpC

Thus far, we observed that perturbation of MrpC NAR led to uncoordinated behavior in which some cells within mature aggregates exited as metastatic swarms and increased variability in *mrpC* expression within the population. To examine the MrpC levels in individual cells and to correlate this with distinct subpopulations, we set out to generate a fluorescent protein fusion to MrpC. Constructs attempting to generate MrpC with fluorescent proteins or small fluorescent tags fused to either the amino- or carboxy-terminus and expressed either from the endogenous *mrpC* locus or from exogenous sites, resulted in strong developmental phenotypes and/or partial release of MrpC from the fusion proteins (data not shown). One interesting exception was a strain bearing a carboxy-terminal mNeonGreen fusion to MrpC expressed from the native *mrpC* promoter in the Δ*mrpC* background (Δ*mrpC attB*::P_WT_-*mrpC-mNeonGreen*; hereafter termed *mrpC*-*mNG*). Surprisingly, this strain exhibited a similar developmental phenotype to that of the P_MUT_-*mrpC*: aggregation onset in the *mrpC*-*mNG* strain occurred 2.5 h earlier than in the parent P_WT_-*mrpC* strain, and metastatic swarms were observed (Fig. S5A, Movie 3). These results strongly suggested that fusion of mNeonGreen to the C-terminus of MrpC was interfering in efficient NAR, likely because the mNG fusion interferes with cooperative multimeric MrpC interactions required for efficient exclusion of MrpB from the *mrpC* promoter region (21). Examination of developing *mrpC*-*mNG* cells by fluorescence microscopy demonstrated that fluorescence was detected primarily in the center of the cell (Fig. S5C) likely colocalized with the nucleoid, consistent with the role of MrpC as a global transcriptional regulator. A similar localization was also observed by anti-MrpC immunofluorescence of harvested developing cells (V. Bhardwaj and P. I. Higgs, unpublished results). Finally, immunoblot analyses of the developmental *mrpC*-*mNG* lysates demonstrated no untagged MrpC could be by detected by polyclonal anti-MrpC immunoblot, indicating MrpC-mNG (predicted molecular mass 54 kDa) was the sole version of MrpC in this strain (Fig. S5B). However, anti-mNG antibodies detected bands corresponding to the 54 kDa MrpC-mNG fusion protein and a 27 kDa mNG monomer (Fig. S5B). We speculate mNeonGreen was released as a result of normal MrpC turnover (23), and was likely detected as diffuse signal which was not localized over the nucleoid. Regardless, mNeonGreen fluorescence indicated the level of MrpC that was (at one point) produced. Therefore, we took advantage of the *mrpC*-*mNG* strain to examine the relative accumulation of MrpC in developing cells in the pre-aggregation, metastatic swarm, sporulating fruiting body, and peripheral rod populations *in situ*. For this purpose, the *mrpC*-*mNG* strain was induced to develop for 24, 30, or 48 hours and stained with a membrane dye (FM4-64) for 60 min prior to imaging by CLSM. At the pre-aggregation stage (24 h post-starvation), we observed randomly aligned rod-shaped cells with a mean per cell mNeonGreen fluorescence of 75 ± 22 arbitrary units (AU) (Fig. 3A-C). By 30 hours, symmetric round aggregates (121 ± 21 AU mean cell fluorescence) and peripheral rods (55 ± 21 AU mean cell fluorescence) were detected. At 48 hours, metastatic swarms were observed, which contained single cell fluorescence values (71 ± 24 AU) between those observed for cells in the residual fruiting bodies (103 ± 31) and peripheral rods (36 ± 23) (Fig. 3A-C). Strikingly, the metastatic swarms consisted almost exclusively of rods aligned with their long axis perpendicular to the moving front of the swarm, whereas the fruiting bodies consisted of spherical shaped spores and rods aligned tangential to the edge of the fruiting body (Fig S5D). At both 30 and 48 hours, the peripheral rods were randomly aligned (Fig S5D). These observations suggest that metastatic swarms arise from cells locked into an intermediate MrpC level that stimulates cell movement, whereas cells that attained a higher level of MrpC reduced cell movement and differentiated into non-motile spores forming immobile fruiting bodies.

**Figure 3.**
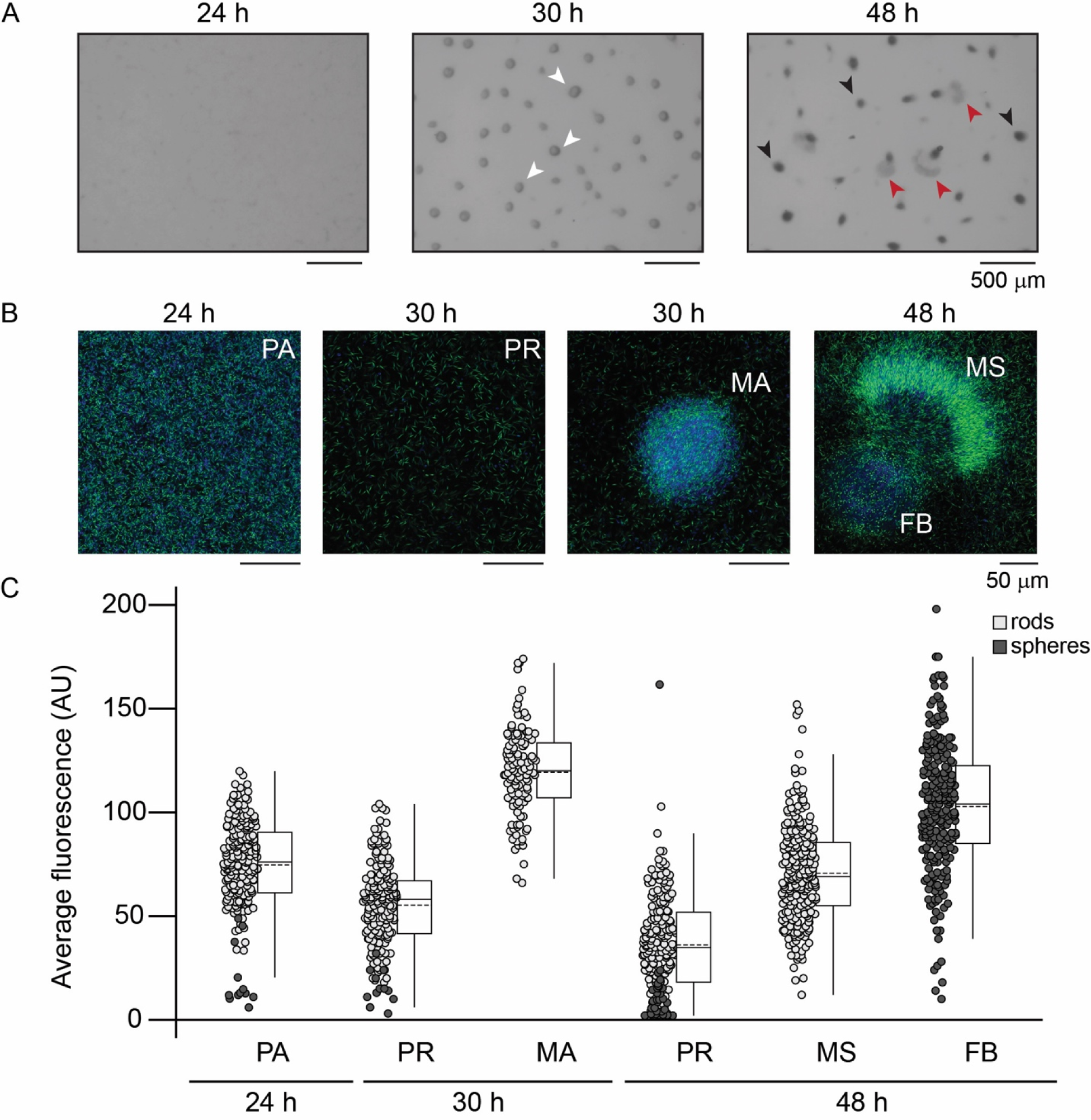
Metastatic swarms correlate with intermediate MrpC production. (A) Developmental phenotype images of the Δ*mrpC* strain expressing *mrpC-mNG* from the wild type *mrpC* promoter integrated at an exogenous locus (P_WT_-*mrpC-mNG*). (B) Fluorescent imaging of P_WT_-*mrpC-mNG* cells in the indicated stages. Development was induced under submerged culture conditions for the indicated times, treated with FM4-64 membrane stain, and imaged by confocal microscopy. Fluorescence captured from the membrane strain and mNeonGreen is colored blue and green, respectively. A and B: Pre-aggregation cells (PA), mature aggregates (MA, white arrows), metastatic swarms (MS, red arrows), and fruiting bodies (FB, black arrows) are indicated. (C) Average mNeonGreen fluorescence recorded from individual cells in each population. Regions of interest (ROIs) were identified based on the membrane stain and the average mNeonGreen fluorescence within the ROI was measured. Results from two independent biological replicates are shown. n = 200 cells for PA, 30 h PR and 48h PR populations. n = 240 cells for 30 h MA, 48 h FB, and 48 h MS populations. Solid line: median; dashed line: mean; light gray circles: rod shaped cells; dark gray circles: spherical cells. Circles in the pre-aggregate and peripheral rod populations likely corresponded to end-on cells, rather that spores.

## Discussion

In this study, we have investigated the role of a native negative autoregulation network motif that is wired into MrpC, a major transcriptional regulator that directs the *M. xanthus* developmental program. MrpC NAR was perturbed by disrupting two MrpC binding sites within the *mrpC* promoter. By expressing *mrpC* from the rewired promoter, we were able to examine the phenotypic consequences of disrupting the NAR motif during *M. xanthus* development. We show that perturbation of MrpC NAR causes cells to produce aggregation centers earlier than the parent (Fig. 1B and C) likely because MrpC accumulated earlier in this strain (Fig. 1D). It has been demonstrated previously that premature MrpC accumulation in *esp* mutants results in early aggregation onset but also in early and increased sporulation efficiency (31) (23). In contrast, the perturbed NAR mutant displayed drastically reduced sporulation efficiency because some cells abruptly exited the aggregation centers and remained locked in a swarming state (Figs. 1E and S2). This phenomenon can be attributed to increased variability in *mrpC* expression (Fig. 2) rather than increased levels *per se*, because an *esp* mutant accumulates MrpC at similar levels and timing to the P_MUT_-*mrpC* strain, but it does not produce metastatic swarms (data not shown). Increased variability in *mrpC* expression likely produces a broad distribution of MrpC such that the population simultaneously contains a mixture of cells that have reached the aggregation and sporulation commitment thresholds (Fig. 4). For example, at 30 h of development, mature aggregates contained cells with levels of MrpC at the upper end of the distribution (Fig. 3), while cells at the lower end have not yet aggregated into these centers. By 48 hours, some of these cells had already reached the level of MrpC required to commit to sporulation and thus remained in stationary fruiting bodies (Fig. 3). Those at the lower end are in the aggregation phase and collectively exited the aggregation center as metastatic swarms. With appropriate NAR, MrpC levels are more homogeneous within a given time frame, and the population would collectively quickly transition from aggregation to sporulation (Fig. 4).

**Figure 4.**
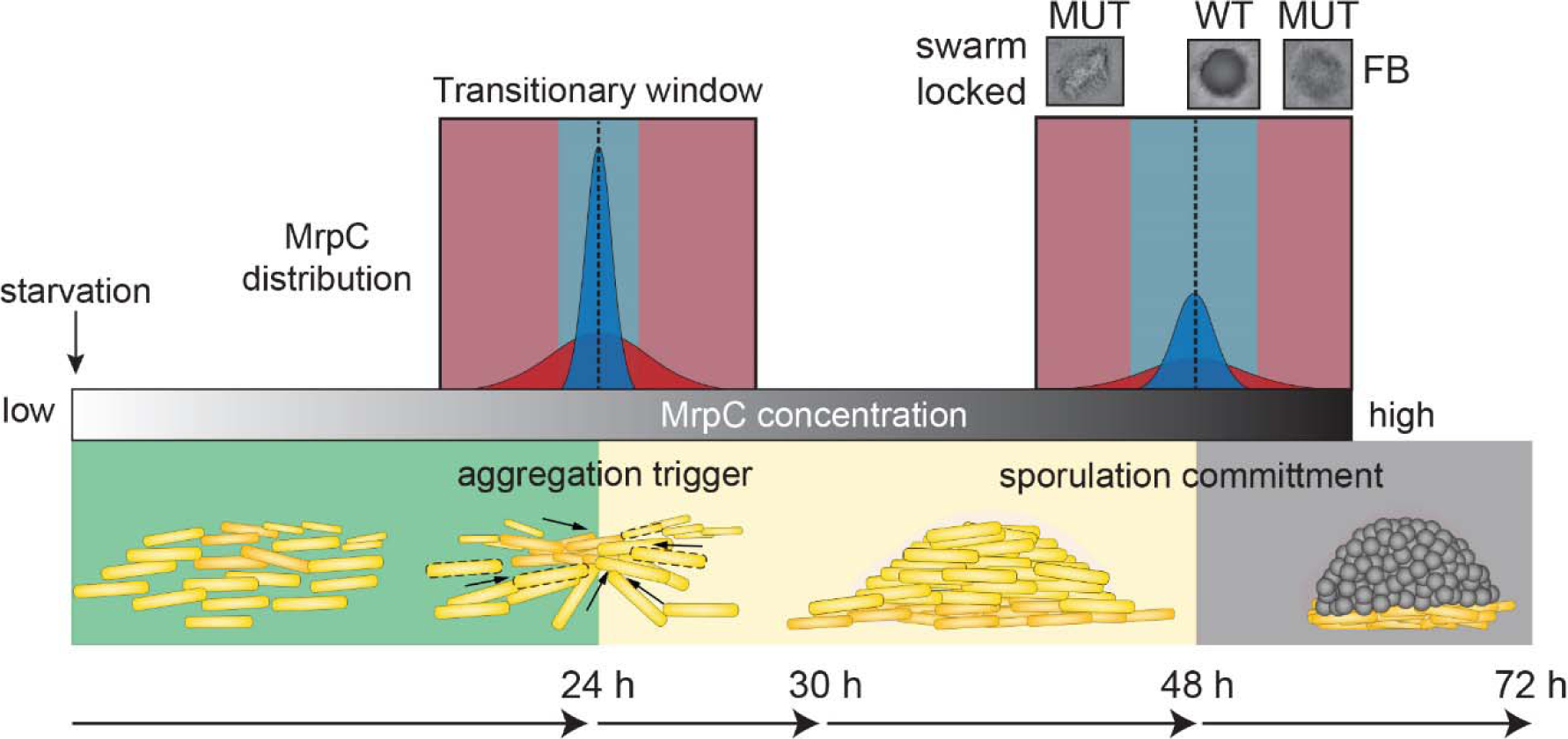
Model for role of NAR in synchronized progression through developmental phases. After onset of starvation, MrpC levels (shaded gray bar) steadily increase. At specific threshold concentrations (dotted lines) aggregation and sporulation commitment are triggered. The duration of pre-aggregation, aggregation and sporulation phases is indicated by green, yellow and grey rectangles, respectively. With appropriate NAR, the distribution of MrpC is constrained (dark blue area), which promotes rapid and synchronized transition of the population to aggregation and then sporulation phases (light blue transitory windows); mature stationary fruiting bodies are produced (WT FB). Perturbation of NAR produces a wider distribution in MrpC concentration (dark red area), which lengthens the time needed for the entire population to transition to the next phase (light red transitory windows). Cells with higher MrpC levels transition to spores in visible fruiting bodies (MUT FB), and those with lower MrpC levels extend the aggregation phase and exit the aggregate as a visible metastatic swarm (MUT swarm locked). See text for details. Schematic of wild type development shown below for comparison. Yellow rods: developing cells; grey circle: spores.

But why did developmental swarms observed in the P_MUT_-*mrpC* strain remain locked in the swarm state? One possibility is that cells inside an aggregation center are subject to an additional positive feedback loop, which is necessary to reinforce commitment to sporulation, *e.g.* C-signaling, a contact-dependent signaling system(32). In cells that left the aggregate, the positive feedback loop would be broken, prohibiting transition to the sporulation phase. A second possibility is that MrpC levels in the developmental swarms continually oscillate between the aggregation and sporulation thresholds. It is known that oscillations can be generated through an additional network motif comprised of a composite negative feedback loop (33). Interestingly, such a loop can be identified for MrpC: MrpC induces expression of *esp* (A. Schramm and P. I. Higgs, unpublished results), and Esp induces degradation of MrpC (23). MrpB induces expression of *mrpC* and MrpC positively regulates expression of *mrpB* (16)(C. Matacyznski and P.I. Higgs, unpublished results).

During development in *M*. *xanthus*, groups of cells organize into a defined pattern (i.e. fruiting bodies) for a designated function (protection and dispersal), making the developing population akin to a specialized bacterial tissue. While multicellular tissue formation in *M*. *xanthus* is relatively simple compared to that of higher eukaryotic organisms, many of the same basic regulatory principles still apply. Cells must synchronously progress through development in a defined temporal order in order to produce a functional structure. This principle is exemplified by the regulation of gastrulation in *Drosophila melanogaster* embryogenesis. Invagination (coordinated cell movement) during gastrulation is largely coordinated by the key regulator, *snail* (34). Expression of *snail* displays a significant degree of homogeneity and synchronicity, which is crucial for its function (35, 36). If synchronicity is perturbed, then multicellular coordination of invagination becomes defective and the severity of the defect strongly correlates with the level of asynchronicity(36). Intriguingly, Snail is proposed to function as a negative autoregulator, which is thought to promote homogeneity of *snail* expression in the population (35). Thus, noise reduction attribute of NAR motifs may therefore be a general strategy for promoting synchronized responses in multicellular developmental systems.

## Materials and Methods

### Bacterial strains, plasmids, and oligonucleotides

The bacterial strains used in this study are listed in Supplemental Table 2. Plasmids were constructed (37) using the oligonucleotide sequences, and construction strategy listed in Supplemental Table 3.

### Growth and Developmental conditions

*Escherichia coli* were grown under standard laboratory conditions in LB media supplemented with 50 μg ml^-1^ of kanamycin and/or 20 μg ml^-1^ of tetracycline, where necessary (38). **M. xanthus** DZ2 strains were grown under vegetative conditions on CYE agar or in broth, as described previously (37); plates were supplemented with 100 μg ml^-1^ of kanamycin and/or oxytetracycline at 10 μg ml^-1^, where necessary.

*M. xanthus* strains were induced to develop under submerged culture conditions (37). Briefly, vegetative cells were diluted to an absorbance at 500 nm (A_550_) of 0.035 in fresh CYE broth, seeded into petri dishes or tissue culture dishes (as indicated in the relevant methods sections) and allowed to grow to a confluent layer for 24 hours at 32 °C. To initiate development, CYE media was removed and replaced with an equivalent volume of MMC buffer [10 mM 4-morpholinepropanesulfonic acid (MOPS) pH 7.6, 4 mM MgSO_4_, 2 mM CaCl_2_], followed by continued incubation at 32 °C for 72-120 hours.

To record static developmental phenotypes, 0.5 ml diluted cells were seeded into 24-well tissue culture plates and imaged at the indicated times with a Leica M80 stereomicroscope.

### Analysis of mCherry fluorescence by plate reader

Submerged culture assays were set up using 0.5 mL diluted cells seeded into each well of 24-well tissue culture plates, and population mCherry fluorescence was measured as described previously (21). Briefly, developing cells were harvested at the indicated hours post-starvation, dispersed, and 1/10 volume of each sample was assayed for fluorescence at 580 nm in a Typhoon imager scan. Values plotted are the average and associated standard deviation from two independent biological replicates.

### Sporulation assay

Developmental sporulation efficiency was determined as described previously from 0.5 mL diluted cells developed in triplicate in 24-well tissue culture plates under submerged culture conditions (37). Briefly, heat (50°C for 60 min)- and sonication (60 pulses 30% output)-resistant spores were enumerated in a Helber counting chamber. Sporulation efficiency was calculated as percent of wild type spores at 72 or 120 hours as indicated. Values reported are the average and associated standard deviation from triplicate independent biological experiments. Chemical-induced sporulation was triggered by addition of glycerol to 0.5 M to vegetatively growing cells in CYE broth with shaking incubation for 24 h at 32°C (39). Spores were isolated and enumerated as indicated above.

### Developmental video analysis

*M. xanthus* strains were induced to develop under submerged culture using 0.15 mL diluted cells per well in 96-well tissue culture plates. After induction of starvation, plates were incubated in a Tecan Spark10M plate reader preheated to 32°C (26). The center of each well was imaged every 30 min from 0 – 72 h post-starvation and images assembled into movies (6 fps) in ImageJ (40). For each movie, onset of aggregation (Agg_ONSET_), maximum aggregates (Agg_MAX_) and final fruiting bodies (Agg_FINAL_) were enumerated as described previously (26). The percent of aggregates that transitioned to stationary fruiting bodies was calculated as [Agg_FINAL_/Agg_MAX_]. The number of aggregates that were mobile (Agg_MOBILE_) in each movie was recorded and percent Agg_MOBILE_ was calculated as [Agg_MOBILE_ / Agg_FINAL_]. For each movie, the first frame (Mobility_I_) and final frame (Mobility_F_) in which aggregates were mobile was recorded and mobility duration was calculated as [(Mobility_F_ - Mobility_I_) × 0.5 h/frame]. Mobility delay was calculated as [Mobility_I_ – Agg_ONSET_]. Data were compiled from three biological replicates that contained five technical replicates per strain.

### Neural network training and analysis

For tracking aggregate and swarm mobility, DeepLabCut deep convolution neural network was used (29). To train DeepLabCut, 769 total frames were extracted from 29 developmental movies that contained aggregates and/or metastatic swarms, which were manually labeled in every frame in which they were present. Using the labeled frames, the DeepLabCut neural network was trained using a 50-layer residual network (30) on Google Colaboratory (hardware accelerator: GPU) for 340,000 iterations (p-cutoff =0.1). The trained neural network possessed a training and test error of 1.62 and of 6.66 pixels, respectively. To track movement of aggregates and swarms, 15 videos for each strain (3 biological replicates each with 5 technical replicates) from 25 – 72 h of development were analyzed with the trained neural network. The predicted labels (likelihood > p-cutoff) for each video were then manually processed and any spurious labels were removed. To track mobility of individual swarms and aggregates, videos were cropped to contain only a single aggregate and/or swarm. Swarms and aggregates were only analyzed for mobility if they remained within the frame for the entirety of the recording. To track mobility in an entire well, developmental videos (640 × 510 pixels) were initially cropped into twenty smaller videos (160 × 102 pixels) with ImageJ (40). Each cropped video was analyzed with the trained neural network as stated previously. Labels from individual videos were then manually stitched together. Displacement between two time points with coordinates (x_i_, y_i_) and (x_f_, y_i_) was calculated as [((x_f_ - x_i_)^2^ + (y_f_ - y_i_)^2^)^-2^ × (1661.5 μm/1280 pixels)]. Total swarm displacement was calculated as the sum of all displacements across all time points. Speed of mobility between two time points (t_i_ and t_f_) was calculated as [displacement/((t_f_ – t_i_) × 60 min/h)].

### Confocal microscopy

For analysis of single cell *mrpC* expression, *M. xanthus* strains bearing P_van_-*mNeonGreen* and either P_WT_-*mCherry* (PH1373) or P_MUT_-*mCherry* (PH1374) were diluted 1:19 with an unlabeled wild type strain (DZ2) and induced to develop under submerged culture conditions using 2.1 mL diluted cells to seed ibiTreated μ-dishes^35mm, high^ (Ibidi). Developing cultures were imaged using a Leica TCS SP8 inverted confocal microscope with a 63x objective. Brightfield images were taken with a gain of 230 V and 0.0% offset. mNeonGreen fluorescence was examined using a 488 nm wavelength laser (5% power) for excitation, a 500 – 540 nm emission spectra, 800 V gain, and 0.0% offset. mCherry fluorescence was examined using a 552 nm wavelength laser (5% power) for excitation, a 585 – 630 nm emission spectra, 650 V gain, and 0.0% offset. For each replicate at 24 h pre-aggregating cells and 48 h peripheral rods, five images were taken from throughout the plate for each strain (line average: 8, resolution: 1024 × 1024). For analysis of the 48 h fruiting bodies, z-stacks of three fruiting bodies were taken for each strain. Each fruiting body was imaged from the base to the interior of the fruiting body (21-37 images) (step size: 1 μm, line average: 4, resolution: 1024 ×1024). Data were compiled from three biological replicates.

Images were subsequently analyzed in ImageJ (40). For 48 h aggregated populations, images were initially cropped to include only the fruiting body. ROIs were created by thresholding the images from the mNeonGreen channel to contain only pixels that were above the intensity of the unlabeled background strain (pixel threshold: 45-255), then analyzing particles with area > 0.5 μm^2^ and circularity of 0.0 – 1.0. ROIs were then transferred to the fluorescent images and the integrated density was measured. The red-green (RG) ratios were plotted and points identified as outliers by Grubb’s test (p < 0.5) were removed. The coefficient of variation (CV) for each biological replicated was calculated by dividing the standard deviation by the mean.

For analysis of MrpC-mNeonGreen production in developing cells, *M. xanthus* strain PH1375 was induced to develop as above. Prior to imaging, FM4-64 was added to 5 μg ml^-1^ final concentration and incubated at 32°C for 60 min. At the designated time points, developing cultures were imaged as above, except mNeonGreen fluorescence was recorded with 500 V gain. FM4-64 fluorescence was detected using 722 – 762 nm emission spectra and 650V gain. For analysis of peripheral rod populations, five images for each replicate were taken throughout the plate (line average: 8, resolution: 1024 × 1024) and ROIs (identified based on membrane stain) were drawn for 20 rod shaped cells and 20 circular cells. For analysis of aggregated cell populations, 1 μm z-stacks of three fruiting bodies or metastatic swarms were recorded. One layer in the z-stack was randomly chosen and ROIs were drawn around 40 circular cells within the fruiting body or 40 rod-shaped cells in the swarms. The mean fluorescence for each ROI in the mNeonGreen channel was measured and plotted.

To account for the relative proportion of spheres and rods in each peripheral rod population, the number of rod-shaped and circular cells were counted in 30.84 × 30.84 pixel ROI (24 hours) or 60.07 × 60.07 pixel ROI (30 and 48 hours) and a proportional equivalent of random cells or spheres were chosen for plotting. All images were analyzed with the Leica Application Suite × (LasX) histogram tool.

### Cell lysate preparation and immunoblot analysis

Cell lysates were generated from strains developed under submerged culture using 16 mL diluted cells seeded in 100-mm petri dishes. At the indicated time points, the overlay buffer was removed, the cell layer was resuspended in 1.5 ml ice-cold MMC. Cells were pelleted (17,000 × *g*, 5 min, 4°C), and pellets were stored at −20°C until further use. 1 ml 13% ice-cold trichloroacetic acid (TCA) and one scoop of 0.1 mm zirconia/silca beads was added to each pellet and subject bead beating with a FastPrep-24 tissue homogenizer (MP Biomedical) at 6.5 m/s for 45 sec at 4°C, six times with 2 min incubation on ice between rounds. Samples were then incubated on ice for at least 15 min. Protein was pelleted as above, washed with 1.0 ml ice-cold Tris buffer [100 mM Tris-HCl (pH 8.0)], then resuspended in 50 μl Tris buffer and 150 μl clear LSB (23), heated at 95°C for 5 min, then stored at −20°C until further use. Protein concentration was determined by Pierce BCA protein assay kit (Thermo Fisher Scientific), samples were diluted to 0.87 μg μl^-1^ in 2 × LSB, and 10 μg protein resolved by denaturing polyacrylamide (10%) gel electrophoresis and transferred (semi-dry) to polyvinylidene difluoride (PVDF) membrane. Membranes were probed with rabbit polyclonal anti-MrpC (1:1,000) and anti-mNeonGreen (1:1000) (Cell Signaling Technologies) and anti-rabbit IgG secondary antibodies conjugated to horseradish peroxidase (HRP) at 1:20,000 or 1:5000, respectively. Signal was detected with enhanced chemiluminescence substrate followed by exposure to autoradiography film.

## Acknowledgments

The authors gratefully acknowledge Justin Kenney for conversations about deep convolution neural network analyses and members of the Higgs and Schrader lab groups for helpful discussions. This work was funded by a National Science Foundation grant IOS-1651921.

