## Supplemental data for "A negative autoregulation network motif is required for synchronized *Myxococcus xanthus* development"

* Penelope I. Higgs

**This PDF file includes:**

Figures S1 to S5

Tables S1 to S3

Legends for Movies S1 to S3

SI References

**Other supplementary materials for this manuscript include the following:**

Movies S1 to S3

Supplementary Figures

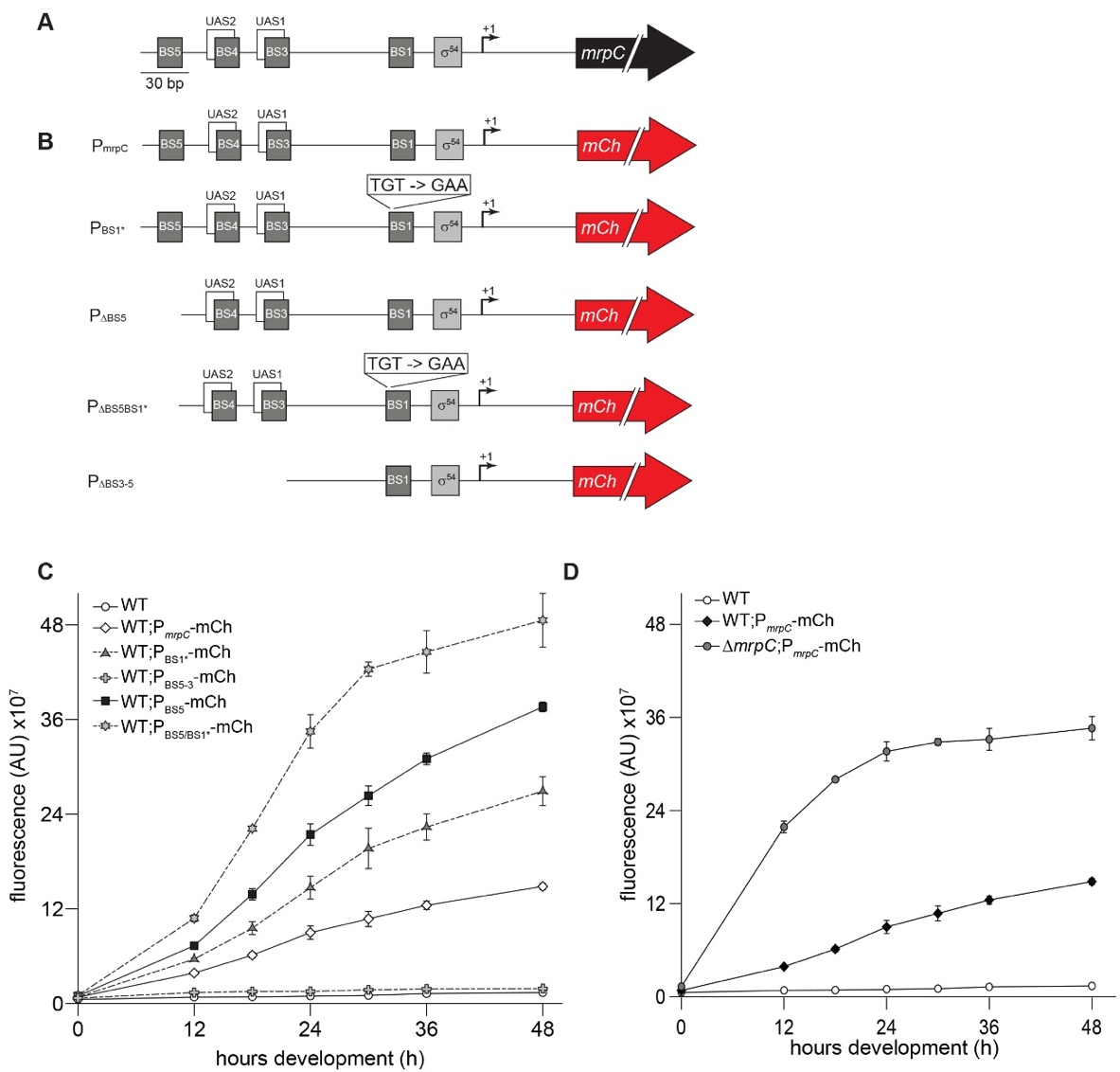

**Figure S1.** MrpC binding sites 1 and 5 each contribute independently to MrpC negative autoregulation. (A) Schematic of the *mrpC* promoter region. BS: Sequences to which MrpC directly binds; UAS: upstream activating sequences to which MrpB directly binds; σ^54^: putative sigma^54^-dependent promoter; +1: transcriptional start site; black arrow: *mrpC* gene. (B) Schematic of the reporter constructs used in (C). P_mrpC_-mCh: the wild type *mrpC* promoter region in (A) was fused to the mCherry fluorescence reporter gene (red arrow). Reporters with mutations/deletions in the *mrpC* promoter are shown below. Disruption of MrpC BS1 was generated by the indicated substitution previously shown to abolish MrpC binding (1). (C) *mrpC* expression as reported from the constructs in (B). Reporters were integrated into the *attB* Mx8 phage attachment site in wild-type cells and mCherry fluorescence was recorded at the indicated hours of development under submerged culture. AU: arbitrary units. (D) Relative *mrpC* expression from the wild type *mrpC* reporter (P*_mrpC_*-mCh) in the wild type or Δ*mrpC* backgrounds. Strains were induced to develop as in C.

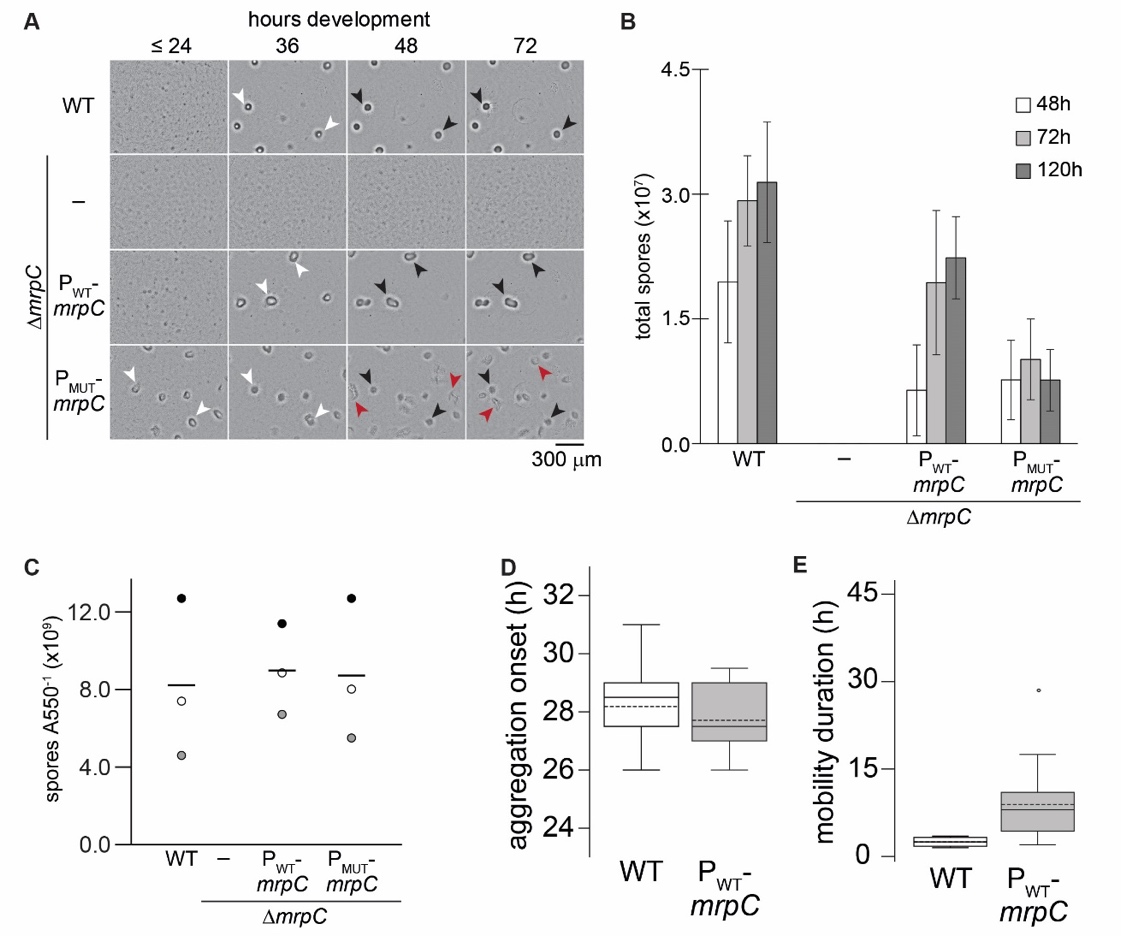

**Figure S2.** Developmental characteristics of wild-type, Δ*mrpC*, P_WT_-*mrpC* and P_MUT_-*mrpC* strains. (A) Comparison of developmental phenotypes of Δ*mrpC* cells expressing *mrpC* from its wild type- (P_WT_-*mrpC*) or P_BS5/BS1-_ (P_MUT_-*mrpC*) promoters (from Fig. 1) to the wild-type and Δ*mrpC* strain. White arrows: aggregates of ~10^5^ cells; black arrows: immobile fruiting bodies; red arrows: metastatic swarms. (B) Spores produced during development under submerged culture. The indicated strains were induced to develop under submerged culture in 24-well tissue culture plates. Cells were harvested at the indicated hours and heat- and sonication-resistant spores were enumerated with a cell counting chamber. Average and associated standard deviations from three independent biological replicates are plotted. (C) Spores produced during chemical induction by the indicated strains. Cells growing in nutrient-rich broth culture were induced to differentiate by addition of glycerol to 0.5M. Cells were harvested after 24 hours and heat- and sonication-resistant spores were enumerated with a cell counting chamber. Spore numbers were normalized to pre-induction cell number (A550). Data plotted from three independent biological replicates are plotted. Replicate 1, 2, and 3 are depicted in white, grey and black circles, respectively. (D-E) Comparison of the wild-type and complemented Δ*mrpC* (Δ*mrpC* *attB*::P_WT_-mrpC) strain phenotypes. Distribution of developmental times at which aggregation centers were first observed (aggregation onset) (D), or durations of time that aggregates traveled (aggregate mobility) (E). Strains were developed under submerged culture in 96 well plates and imaged every 30 min for 72 hours. n=15 from three independent biological replicates. Solid line: median; dashed line: mean.

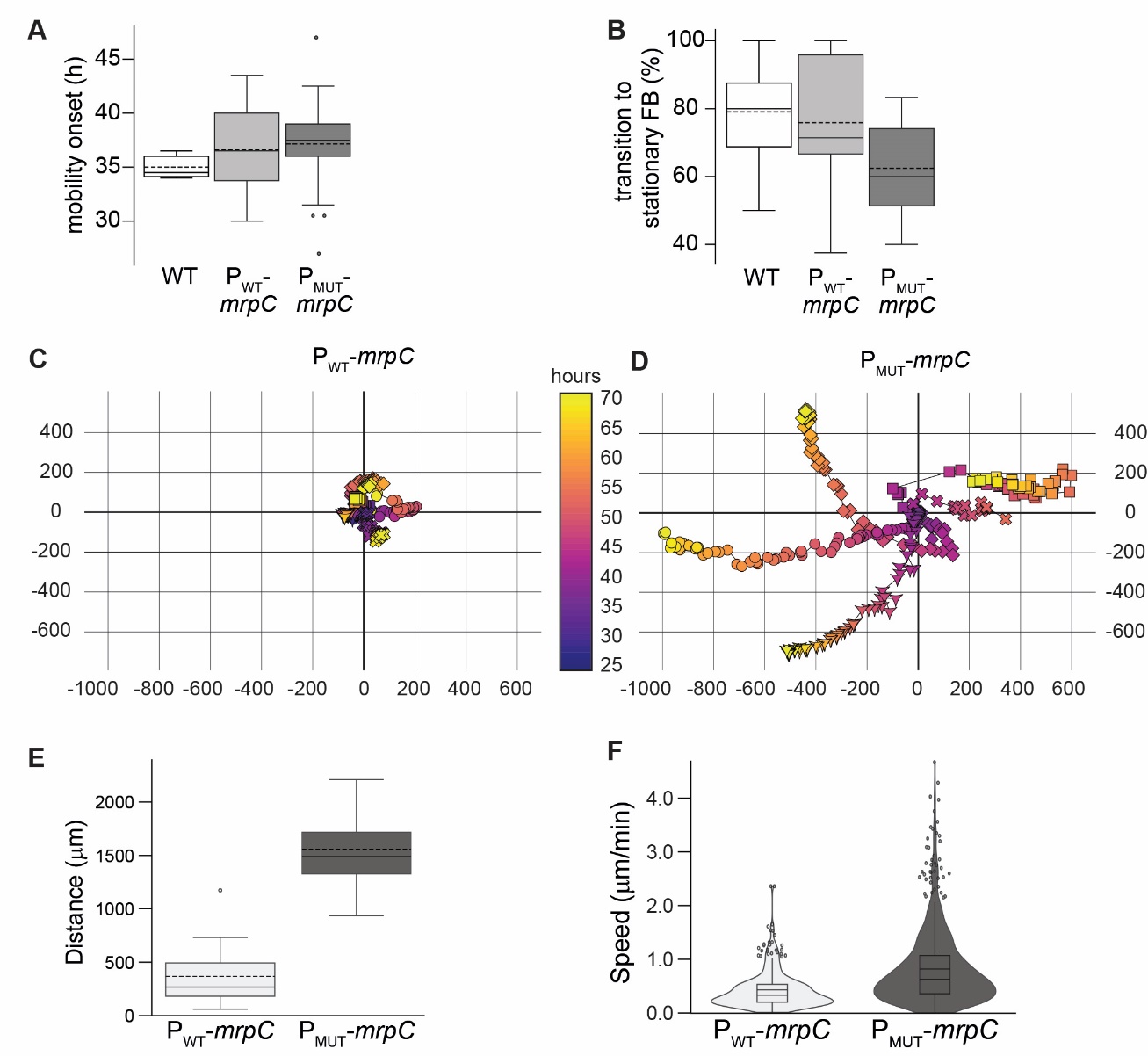

**Figure S3.** Mature aggregate (wild-type, P_WT_-*mrpC*) and metastatic swarm (P_MUT_-*mrpC*) mobility characteristics. (A-B, E-F) Distribution of: developmental times at which mature aggregates or metastatic swarms began to travel (A), percent of initial aggregates that transitioned into mature fruiting bodies (B), and distance travelled (E) and speed (F) of mobile aggregates/metastatic swarms. Strains were developed under submerged culture in 96 well plates and imaged every 30 min for 72 hours. n=15 from three independent biological replicates. Solid line: median; dashed line: mean. (C, D) Track of five different mobile aggregates (P_WT_-*mrpC*)(C) or metastatic swarms (P_MUT_-*mrpC*)(D) from 25 to 72 hours of development. The different aggregates/swarms (indicated by connected circles, squares, diamonds, crosses, and triangles) were set to 0,0 and displacement in the x/y direction was recorded at the times indicated by the heat map. Aggregates/swarms were tracked by trained DeepLabCut neural network.

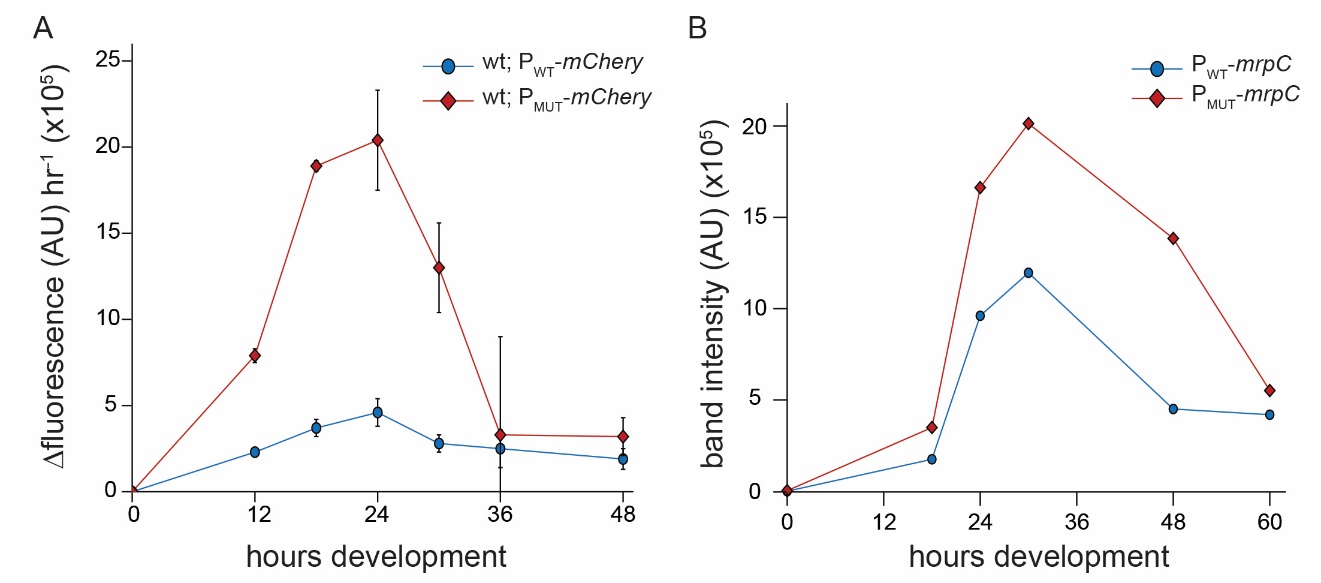

**Figure S4.** Comparison of *mrpC* expression and MrpC protein levels under wild type and perturbed MrpC NAR promoters. (A) mCherry fluorescence from Fig. 1A plotted as the change in mCherry fluorescence normalized to time to account for mCherry stability. (B) Relative intensity of MrpC as detected from anti-MrpC immunoblot from Figure 1D.

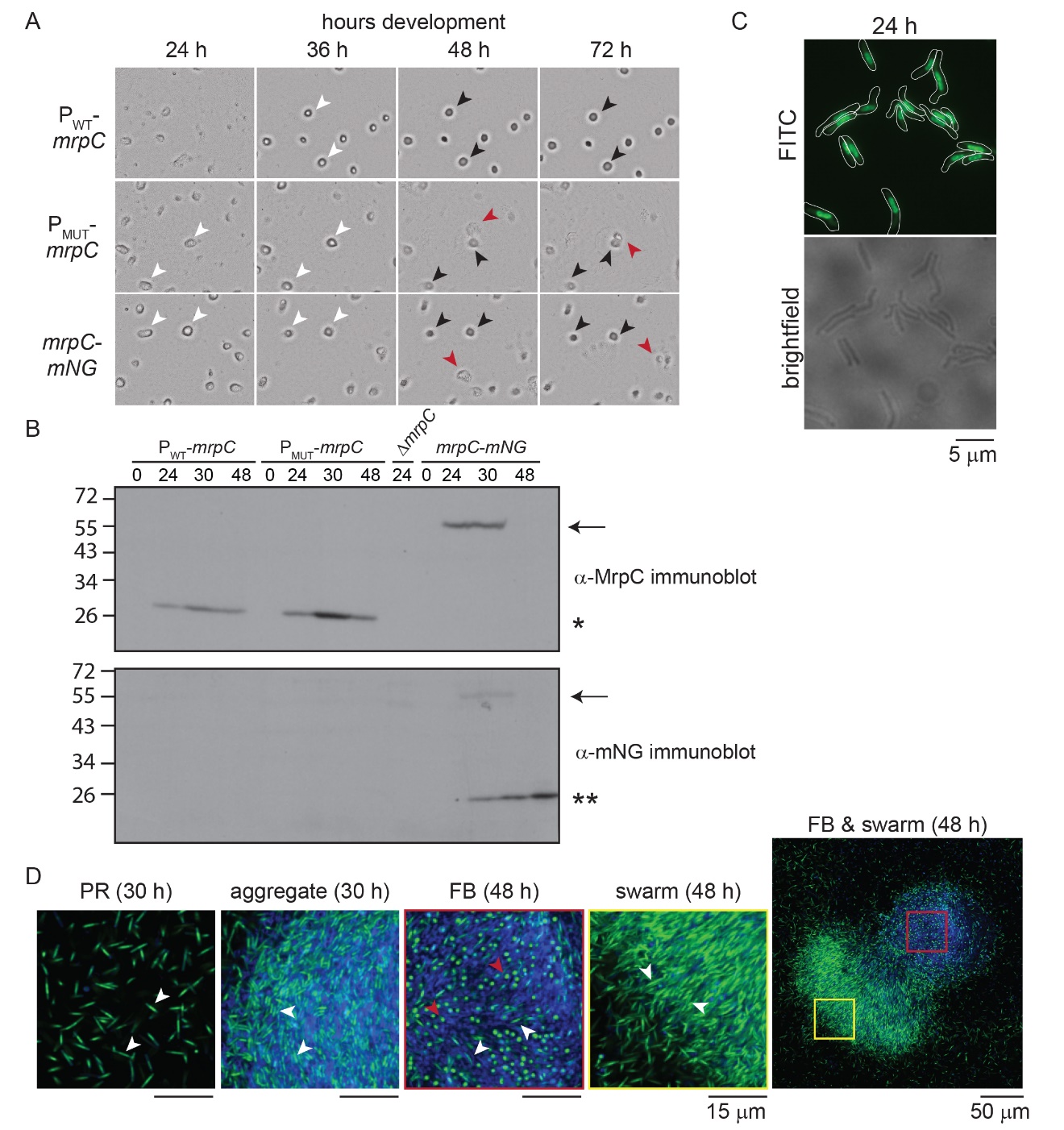

Figure S5. Developmental phenotypes and MrpC protein levels in the *mrpC-mNG* strain. (A) Comparison of developmental phenotypes of Δ*mrpC* cells expressing *mrpC* from its wild type- (P_WT_-*mrpC*) or P_BS5/BS1-_ (P_MUT_-*mrpC*) promoters or *mrpC-mNeonGreeen* from the wild type promoter. Cells were induced to develop under submerged culture in 96 well plates. White arrows: aggregates of ~10^5^ cells; black arrows: immobile fruiting bodies; red arrows: metastatic swarms. (B) Anti-MrpC (top) or anti-mNeonGreen (bottom) immunoblots of cell lysates prepared from the indicated strains developing under submerged culture in 100 mm petri dishes. MrpC, MrpC-mNeonGreen, and truncated mNeonGreen are indicated by single star, arrow, and double star, respectively. No MrpC or mNeonGreen is detected in the 24 hour Δ*mrpC* lysate. (C) MrpC-mNeonGreen localizes to the center of the cell. The *mrpC-mNG* strain was induced to develop under submerged culture for 24 hours and cells were harvested, dispersed, and imaged by epifluorescence microscopy. Top: Fluorescence detected with FITC filters. Bottom: brightfield image. Size bar: 5 μm. (D) Alignment of rods (white arrows) and/or spores (red arrows) and the respective MrpC-mNeonGreen localization in the peripheral rod (PR), mature aggregate (MA), fruiting body (FB) or metastatic swarm (MS) populations. Images outlined in red and yellow correspond to the boxes in the right-most panel showing a metastatic swarm leaving a residual fruiting body. *mrpC-mNG* cells were induced to develop under submerged culture for the indicated times, stained with FM-464 membrane stain, and imaged under a confocal microscope. Fluorescence captured from the membrane strain and mNeonGreen is colored blue and green, respectively.

Table S1. Developmental phenotype characteristics.

| Characteristics^a^ | WT | P_WT_-*mrpC* | P_MUT_-*mrpC* |
| --- | --- | --- | --- |
| Aggregation onset (h) | 28 ± 1 h | 28 ± 1 h | 25 ± 1 h |
| Max aggregates | 8 (IQR: 6 – 9) | 6 (IQR: 5 – 8) | 8 (IQR: 6 – 10) |
| Final aggregates | 6 (IQR: 5 – 8) | 5 (IQR: 4 – 6) | 5 (IQR: 4 – 6) |
| Transition to mature FB (%) | 80 ± 20% | 80 ± 20% | 60 ± 10% |
| Mobile aggregates (%) | 5 ± 10% | 40 ± 30% | 80 ± 20% |
| Mobility onset (h) | 34 ± 2 h | 37 ± 4 h | 37 ± 4 h |
| Mobility delay (h) | 4 ± 3 h | 7 ± 3 h | 10 ± 3 h |
| Mobility duration (h) | 2 ± 1 h | 9 ± 6 h | > 29 h |

^a^see text for descriptions

Table S2. Strain and plasmid used in this study.

| *M. xanthus* strains | | |
| --- | --- | --- |
| Strain | **Genotype** | **Source** |
| DZ2 | Wild type | (2) |
| PH1025 | DZ2 ∆*mrpC* | (3) |
| PH1100 | DZ2 *attB*::pPTM014 (P*_mrpC_*-*mCherry*), Km^R^ | (1) |
| PH1104 | PH1025 *attB*::pPTM014 (P*_mrpC_*-*mCherry*), Km^R^ | (1) |
| PH1161 | DZ2 *attB*::pPTM022 (P*_mrpC_*_(∆BS3-5)_-*mCherry*), Km^R^ | (1) |
| PH1162 | DZ2 *attB*::pPTM024 (P*_mrpC_*_(∆BS5)_-*mCherry*), Km^R^ | (1) |
| PH1370 | DZ2 *attB*::pPTM019 (P*_mrpC_*_(BS1 TGT→GAA )_-*mCherry*), Km^R^ | This study |
| PH1371 | DZ2 *attB*::pPTM029 (P*_mrpC_*_(∆BS5/BS1 TGT→GAA )_-*mCherry*), Km^R^ | This study |
| PH1118 | PH1025 *attB*::pVG114 (P*_mrpC_*-*mrpC*), Km^R^ | (1) |
| PH1372 | DZ2 *attB*::pPTM029 (P*_mrpC_*_(∆BS5/BS1 TGT→GAA )_-*mrpC*), Km^R^ | This study |
| PH1373 | PH1100 1.38-kb::pPTM032 (P_vanillate_-*mNeonGreen*), Km^R^ Tc^R^ | This study |
| PH1374 | PH1371 1.38-kb::pPTM032 (P_vanillate_-*mNeonGreen*), Km^R^ Tc^R^ | This study |
| PH1375 | PH1025 *attB*::pPTM037 (P*_mrpC_*-*mrpC*-(gly_4_-ser)_2_-*mNeonGreen*), Km^R^ | This study |
| *E. coli* strains | | |
| Strain | **Genotype** | **Source** |
| TOP10 | F^—^ *endA1 recA1 galE15 galK16 nupG rpsL* ∆*lacX74* Φ80*lacZ*∆M15 *araD139* ∆(ara, leu)7697 *mcrA* ∆(*mrr-hsdRMS*-*mcrBC*) λ^—^ | Invitrogen |
| Plasmid | **Genotype** | **Source** |
| pPTM014 | pSL8 P*_mrpC_*-*mCherry*, Km^R^ | (1) |
| pPTM019 | pSL8 P*_mrpC_*_(BS1 TGT→GAA)_-*mCherry*, Km^R^ | This study |
| pPTM022 | pSL8 P*_mrpC_*_(∆BS3-5)_-*mCherry*, Km^R^ | (1) |
| pPTM024 | pSL8 P*_mrpC_*_(∆BS5)_-*mCherry*, Km^R^ | (1) |
| pPTM029 | pSL8 P*_mrpC_*_(∆BS5/BS1 TGT→GAA)_-*mCherry*, Km^R^ | This study |
| pPTM030 | pFM18 P*_mrpC_*_(∆BS5/BS1 TGT→GAA)_-*mrpC*, Km^R^ | This study |
| pPTM032 | pMR3629 P_vanillate_-*mNeonGreen*, Tc^R^ | This study |
| pPTM037 | pFM18 P*_mrpC_*-*mrpC*-(gly_4_-ser)_2_-*mNeonGreen*, Km^R^ | This study |
| pVG114 | pSL8 P*_mrpC_*-*mrpC*, Km^R^ | (1) |

**Table S3.** **Oligonucleotide sequences and construction strategies of plasmids used in this study.**

| Plasmid^a^ | Description^b^ | Primer  type^c^ | Oligo name | Sequence (5'-3')^d^ |
| --- | --- | --- | --- | --- |
| pPM019 | P*_mrpC_*_(BS1 TGT→GAA)_-*mCherry* | A | oPH500 | GACCCGGGTCTTGCACAGAGCCAGAG |
|  |  | B | oPH1802 | TTGAAATGCCGCCGGGACATTCCC |
|  |  | C | oPH1803 | ATTTCAGGAGGGAATGTCCC |
|  |  | D | oPH503 | GGTCTAGATCACTTGTACAGCTCGTCCATGC |
| pPM029 | P*_mrpC_*_(BS5/BS1 TGT→GAA)_-*mCherry* | A | oPH1885 | GACCCGGGGCCCCCGGTTTGAAC |
|  |  | B | oPH1802 | TTGAAATGCCGCCGGGACATTCCC |
|  |  | C | oPH1803 | ATTTCAGGAGGGAATGTCCC |
|  |  | D | oPH1869 | GGTCTAGATCACTTGTACAGCTCGTCCATGC |
| pPM030 | P*_mrpC_*_(BS5/BS1 TGT→GAA)_-*mrpC* | A | oPH1885 | GACCCGGGGCCCCCGGTTTGAAC |
|  |  | B | oPH1802 | TTGAAATGCCGCCGGGACATTCCC |
|  |  | C | oPH1803 | ATTTCAGGAGGGAATGTCCC |
|  |  | D | oPH947 | CGATAAGCTTCTACTTCTCCTTGCCG |
| pPTM032 | P_vanillate_-*mNeonGreen* | for | oPH1920 | CGATCATATGGTCAGCAAAGGTGAAG |
|  |  | rev | oPH1921 | CGAGAATTCTCACTTGTACAGTTCGTCC |
| pPTM037 | P*_mrpC_*-*mrpC-mNeonGreen* | for1 | oPH1908 | GCAGATCATATGTCTTGCACAGAGCCAGAGC |
|  |  | rev1 | oPH1982 | GGAGCCGCCGCCGCCCTTCTCCTTGCC |
|  |  | for2 | oPH2003 | GGCGGCGGCGGCTCCATGGTCAGCAAAGGTG |
|  |  | rev2 | oPH2004 | TCGAAGCTTTCACTTGTACAGTTCGTCC |

^a^Plasmid name, ^b^Plasmid insert description, ^c^ Primers labeled as A, B, C, or D represent the primers used for the two-step PCR fragment fusion process [described in (4)]. Primers are otherwise labeled as forward (for) or reverse (rev). ^d^Underlined sequences indicate restriction sites used for cloning

Movie S1. Developmental phenotype of the P_WT_-*mrpC* strain. Cells were induced to develop under submerged culture and imaged every 30 minutes from 0 – 72 h (5).

Movie S2. Developmental phenotype of the P_MUT_-*mrpC* strain. Cells were induced to develop under submerged culture and imaged every 30 minutes from 0 – 72 h (5).

Movie S3. Developmental phenotype of the *mrpC*-*mNeonGreen* strain. Cells were induced to develop under submerged culture and imaged every 30 minutes from 0 – 72 h (5).
